# Concussion-related disruptions to hub connectivity in the default mode network are related to symptoms and cognition

**DOI:** 10.1101/2023.03.07.531551

**Authors:** Heather C. Bouchard, Kate L. Higgins, Grace K. Amadon, Julia M. Laing-Young, Arthur Maerlender, Seima Al-Momani, Maital Neta, Cary R. Savage, Douglas H. Schultz

**Affiliations:** Center for Brain, Biology and Behavior, University of Nebraska-Lincoln, Lincoln, NE; Department of Psychology, University of Nebraska-Lincoln, Lincoln, NE; Department of Athletics, University of Nebraska-Lincoln, Lincoln, NE; Department of Neurosurgery, Medical College of Wisconsin, Milwaukee, WI

**Keywords:** sport-related concussion, network hub, mild traumatic brain injury, default mode network, functional connectivity, neuroimaging

## Abstract

Concussions present with a myriad of symptomatic and cognitive concerns; however, the relationship between these functional disruptions and the underlying changes in the brain are not yet well understood. Hubs, or brain regions that are connected to many different functional networks, may be specifically disrupted after concussion. Given the implications in concussion research, we quantified hub disruption within the default mode network (DMN) and between the DMN and other brain networks. We collected resting-state functional magnetic resonance imaging data from collegiate student-athletes (n = 44) at three timepoints: baseline (prior to beginning their athletic season), acute post-injury (approximately 48 hours after a diagnosed concussion), and recovery (after starting return-to-play progression, but prior to returning to contact). We used self-reported symptoms and computerized cognitive assessments collected across similar timepoints to link these functional connectivity changes to clinical outcomes.

Concussion resulted in increased connectivity between regions within the DMN compared to baseline and recovery, and this post-injury connectivity was more positively related to symptoms and more negatively related to visual memory performance compared to baseline and recovery. Further, concussion led to decreased connectivity between DMN hubs and visual network non-hubs relative to baseline and recovery, and this post-injury connectivity was more negatively related to somatic symptoms and more positively related to visual memory performance compared to baseline and recovery. Relationships between functional connectivity, symptoms, and cognition were not significantly different at baseline versus recovery.

These results highlight a unique relationship between self-reported symptoms, visual memory performance and acute functional connectivity changes involving DMN hubs after concussion in athletes. This may provide evidence for a disrupted balance of within- and between-network communication highlighting possible network inefficiencies after concussion. These results aid in our understanding of the pathophysiological disruptions after concussion and inform our understanding of the associations between disruptions in brain connectivity and specific clinical presentations acutely post-injury.

## Introduction

Approximately 3.8 million concussions occur each year within the United States^1^ – approximately 300,000 of which are sports- or recreation-related^2^ – and the incidence of sports-related concussions has been steadily increasing^3,4^. These increases in diagnosis may be due to increased awareness, changes in reporting requirements, changes in sport participation, and better recognition of concussion-related symptoms^5^. Unfortunately, we still have limited understanding of the underlying brain pathophysiology resulting from a concussion and of the relationship between the clinical presentation of concussion and physiological brain changes that have been identified. The clinical presentation of concussions can include a variety of symptoms and changes in cognition, including somatic complaints (i.e., sensitivity to light or noise), cognitive deficits (i.e., difficulty concentrating or acute disruptions in memory), affective dysregulation (i.e., feeling more emotional), and sleep-related concerns (i.e., drowsiness).

The definition of a concussion and criteria for diagnosis can vary between practitioners, organizations, and consensus statements^6^. Typically, diagnoses are based on several clinical signs that represent a functional disturbance in the brain^7^. Evidence on standard clinical brain imaging (most commonly CT or structural MRI) is not currently required for the diagnostic process due to lack of sensitivity to these mild brain related changes^7,8^. However, recent research using more advanced MRI techniques has detected subtle changes in brain structural and functional connectivity, which has increased the understanding of the neurophysiological changes in the brain after concussion, specifically related to changes in axonal damage, inflammation, and cellular homeostasis^9^.

Reviews of the literature on functional MRI have reported both increases and decreases in functional connectivity post-concussion, evident globally and within a range of functional brain networks^10,11^. These inconsistencies have limited interpretation of these findings, which may be related to variability in measurement timelines post-injury. Given the heterogeneity of the literature and heterogeneity in the clinical presentation of concussions, it is unclear how the clinical presentation of a concussion relates to changes identified in brain function. Defining these connections may aid in mechanism-based interventions for concussion management.

Patients with moderate to severe traumatic brain injuries (TBIs) have disruptions in functional brain regions identified as “hubs”^12–15^. Hubs are well-connected regions – evident in nearly all functional networks of the brain – that play an important role in brain organization as well as cognitive and behavioral functioning^15–17^. Specifically, hubs are involved in reducing the metabolic cost associated with communication across the brain and allow for greater global efficiency of the brain’s communication^19,20^. Therefore, when hubs are disrupted through pathological processes, there is a disproportionate decrease in network efficiency compared to disruption of non-hubs. For example, lesions to regions identified as hubs can lead to long-term cognitive deficits^16^,, and more subtle injuries, such as a concussion, may also result in a disruption to hubs^22–25^. However, this research on concussions (typically classified as mild TBIs) is limited and only includes imaging data collected post-injury.

The default mode network (DMN) is thought to be central to concussion-related pathophysiology^11,27,28^. It plays a role in a wide variety of cognitive processes, including cognitive and emotional processing, monitoring of the environment, shifting between cognitive tasks, self-referential processing (i.e. remembering the past, planning the future, and self-reflection), and day dreaming^29–31^, many of which are impacted following a concussion. As a result, the DMN is one of the most investigated functional brain networks within the concussion literature^11^. Reported changes to DMN functional connectivity following concussion have been variable, including findings of both hyperconnectivity and hypoconnectivity after injury^11,32^. However, studies are heterogeneous in their neuroimaging preprocessing steps, analytical models, mechanism of injury, ages of participants, and timing of post-injury assessments. In addition, these studies did not include a measure of functional connectivity prior to injury. Variability exists in baseline functional connectivity across people due to distinct individual-specific features of the brain’s cortical organization. The previous inability to capture these intraindividual effects of concussion on the DMN may contribute to the heterogeneity of the literature.

An additional confound in understanding concussion-related physiological changes, such as functional connectivity disruptions, is the extended longevity of these changes. Often these changes persist past the point of recovery as defined by traditional clinical measures^33^. Time between concussion diagnosis and return-to-play for collegiate athletes occurs on average in sixteen days^34^. However, functional connectivity differences are still present up to 23 months after athletes returned-to-play^33^. It is unclear why these physiological changes continue after clinical recovery. One explanation may be due to the brain’s ability to compensate. Another explanation may be from comparing effects between groups (i.e., non-injured control group), as interindividual differences in cortical organization may complicate interpretations. Comparing post-injury functional connectivity patterns to an athlete’s own baseline may resolve some heterogeneity in the literature.

This study aimed to explicate concussion-related changes to DMN functional connectivity, focusing on changes within and between DMN hubs, using a baseline comparison method. Collegiate student-athletes’ functional connectivity at baseline (before the start of the season), was compared to connectivity approximately 48 hours after receiving a concussion diagnosis (post-injury), and after beginning their return-to-play progression, but prior to return to contact (recovery). We hypothesized that functional connectivity of hubs *within* the DMN would be disrupted acutely post-injury compared to both baseline and recovery. Similarly, we hypothesized that as symptom reporting and cognitive performance return to baseline levels at recovery, so will functional connectivity differences. Lastly, for between-network functional connectivity, we predicted that functional connectivity *between* DMN hubs and other networks would be disrupted following a concussion and the degree of this disruption would be related to self-reported symptoms and cognitive performance. However, we did not expect that all disruptions *between* the DMN and other networks will return to baseline levels at recovery, suggesting compensatory processes that allow for the appearance of clinical recovery while changes in the brain persist.

## Methods

### Participants

Forty NCAA men’s football and four women’s soccer student-athletes participated in the study. At baseline, participants ranged in age from 17-24 years (*M* = 19.75, *SD* = 1.8). Participants completed an MRI scan during their baseline clinical evaluation prior to the start of their season. However, during the first year of data collection, a subset of participants (15 football; 2 soccer) had already been competing at the university and MRI scans were conducted at the start of that season. Acute post-injury scans were collected within approximately 48 hours of a diagnosed concussion (post-injury) and during their return to play protocol immediately prior to returning to contact (recovery). Recovery time for participants ranged from 3-93 days (*M* = 11, *SD* = 14). Additional demographic information, including sample sizes due to missing clinical data at the three timepoints, can be found in Table 1.

**Table 1.** Sample Demographics.

|  |  |  |  |  |
| --- | --- | --- | --- | --- |
| Age | **Range**<br>17-24 years |  | **Mean (SD)**<br>19.8 (1.8) |  |
| Recovery Time | 3-93 days |  | 10.5 (13.5) |  |
| Gender | **Women**<br>n = 4 |  | **Men**<br>n = 44 |  |
| Race | **White**<br>n = 22 | **Black**<br>n = 12 | **Biracial**<br>n = 1 | **Unknown**<br>n = 4 |
| Concussion History | **Zero**<br>n = 22 | **One**<br>n = 16 | **Two**<br>n = 4 | **Three**<br>n = 1 |
| Participants with clinical data | **Baseline**<br>n = 44 |  | **Post-Injury**<br>n = 40 | **Recovery**<br>n = 31 |
| Days between MRI scans | **Baseline-Post-Injury**<br>**Mean (SD)**<br>265.1 (254.1) |  | **Post-Injury-Recovery**<br>**Mean (SD)**<br>10.5 (13.7) | **Baseline-Recovery**<br>**Mean (SD)**<br>275.7 (245.3) |
|  | **Minimum – Maximum**<br>19 days – 1028 days |  | **Minimum – Maximum**<br>6 days – 93 days | **Minimum – Maximum**<br>27 days – 1034 days |

Clinical data regarding concussion diagnosis, management, and clearance for return-to-play, including symptom reporting and cognitive performance, were collected in accordance with the Department of Athletics’ concussion protocol at University of Nebraska-Lincoln (UNL). Concussions were diagnosed, managed, and cleared by the concussion management team in the Department of Athletics using a standardized protocol. Diagnoses were provided by the same clinicians and return-to-play criteria included return to baseline symptom levels and clinical judgment. Most concussions were sport-related (n = 41), while three were not sport-related.

Most participants (n = 27) completed their clinical baseline assessments an average of 21 days (range 0 – 65 days) prior to their baseline MRI scan. A subset during the first year of data collection experienced a longer duration between their clinical baseline assessments and MRI scan with an average of 1.8 years (639 days, range 113 days – 3.1 years). This was due to clinical baseline assessments being collected prior to the start of the neuroimaging study. The clinical baseline measures represent normative function that is thought to be relatively stable over time. However, we included time in the regression models as a nuisance variable to account for potential variance. Clinical data, such as symptom reporting and cognitive performance assessed at baseline were not missing for any participants, nor were MRI data at any of the three timepoints (i.e., baseline, post-injury, recovery). However, due to clinical management procedures, clinical data used in this study for fifteen participants was missing at either post-injury or recovery timepoints. All participants signed consent forms approved by the UNL Institutional Review Board to use their MRI scan and clinical data for research purposes.

### Symptom Reports & Cognitive Assessment

Self-reported symptoms and cognitive performance were assessed via Immediate Post-Concussion Assessment and Cognitive Testing (ImPACT)^35^. Self-report of 22 symptoms were collected via the ImPACT Post-Concussion Symptom Scale (PCSS)^36^ at each timepoint, and grouped into four symptom factors: cognitive, somatic, sleep, and affective^37^. ImPACT also creates four composite scores including reaction time, verbal memory, visual memory, and visual-motor speed and was administered at baseline, post-injury, and before clearance to begin return-to-play protocol (recovery) by clinicians in the Department of Athletics. To account for practice effects and regression to the mean, composite scores were converted to regression-based z-scores using a formula derived from a previously published sample who also completed ImPACT across several timepoints^38^.

### MRI Acquisition

Data were collected on a 3T Siemens Skyra scanner housed within the Center for Brain, Biology, and Behavior at UNL. T1-weighted anatomical (TR = 2530 ms, flip angle = 7°, FOV = 256 mm/100% phase, 176 slices, slice thickness = 1 mm) and multiband echo-planar resting-state fMRI (TR/TE = 1000/29.8 ms, flip angle = 60°, FOV = 210 mm/100% phase, slice thickness = 2.5 mm, 51 interleaved slices, multiband accel factor = 3) were acquired. During resting-state fMRI, Framewise Integrated Real-Time MRI Monitoring (FIRMM) was used to assess real-time head motion of participants during their scan to improve data quality^39^. In the case of participant motion, scans were extended until we reached our goal of 30 minutes of low-motion data (framewise displacement < 0.2 mm) or until allocated scan time was over (mean [standard deviation] scan time in minutes for baseline = 28.3 [4.6], post-injury = 29.6 [3.8], recovery = 30.8 [2.6]).

### Resting-state fMRI preprocessing

Data were processed in Matlab (v.R2016b) with a previously established pipeline described elsewhere^40^. Preprocessing occurred with the Talairach atlas^41^ space to register the mean intensity of the blood-oxygen-level-dependent (BOLD) signal with the T1-weighted image. Masks were created of the whole brain, white matter, and ventricles using Freesurfer’s recon-all command^42^ (v5.3). The following preprocessing steps all occurred in Matlab (v.R2016b). This involved slice-time correction, rigid body realignment, and normalization. Six motion estimates and their derivatives^43^ were demeaned and detrended and were included as nuisance regressors along with global signal, white matter signal, and cerebral spinal fluid signal. Frames with high motion (framewise displacement > 0.2 mm) were removed. These scrubbed time points were replaced with interpolated data, using least squares spectral estimation, for the purposes of bandpass filtering but were excluded when calculating functional connectivity^44^. Next, temporal filtering was conducted with a bandpass filter between 0.009 to 0.08 Hz and spatially smoothed with a Gaussian filter of 6 mm. After preprocessing steps were complete, the mean signal from 10mm diameter spheres for each of 264 nodes^45^ was extracted within Matlab (v.R2016b). Matrices representing the correlation (Pearson’s r) between each pair of nodes were calculated and Fisher’s z-transformed.

### Hub identification

Participation coefficient^46^ is a graph theoretic measure used to quantify the uniformity of node distribution within its own network or between other networks. We used participation coefficient to define hubs and non-hubs during participants’ baseline scan in each of the 13 networks described by Power and colleagues^45^ (the “uncertain network” was excluded from this analysis). The conventional participation coefficient measure assumes all networks are the same size, which is not the case in brain network connectivity. Therefore, a *normalized* participation coefficient value was used to reduce the influence of network size^46^. Hubs in each network were identified as having a participation coefficient value in the top 25% of their network, while the remaining nodes in the network were identified as non-hubs. Figure 1 visualizes the average likelihood of a region being selected as a hub for two networks: DMN and visual network. Thresholds of 15% and 35% were also assessed and results are reported in Supplemental Material 1. Figure 2 represents a schematic of the within and between DMN functional connectivity described in following, corresponding sections.

**Figure 1:**
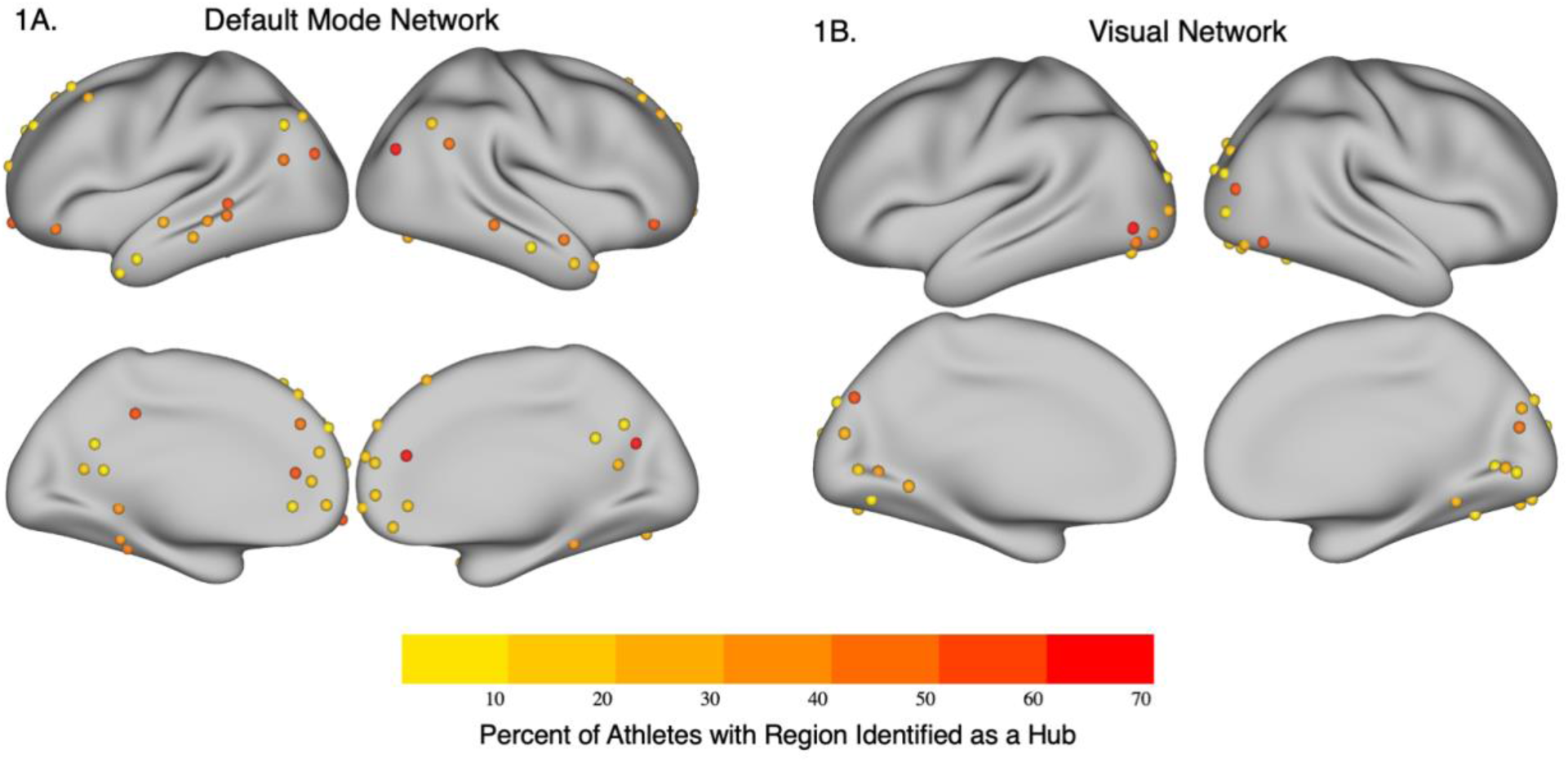
Visualization of Hubs. The average likelihood of a region to be identified as a hub across the 44 student-athletes is visualized across the **(A)** Default Mode Network and **(B)** Visual Network. Regions in red are more likely to be a hub for each participant compared to regions in yellow.

**Figure 2:**
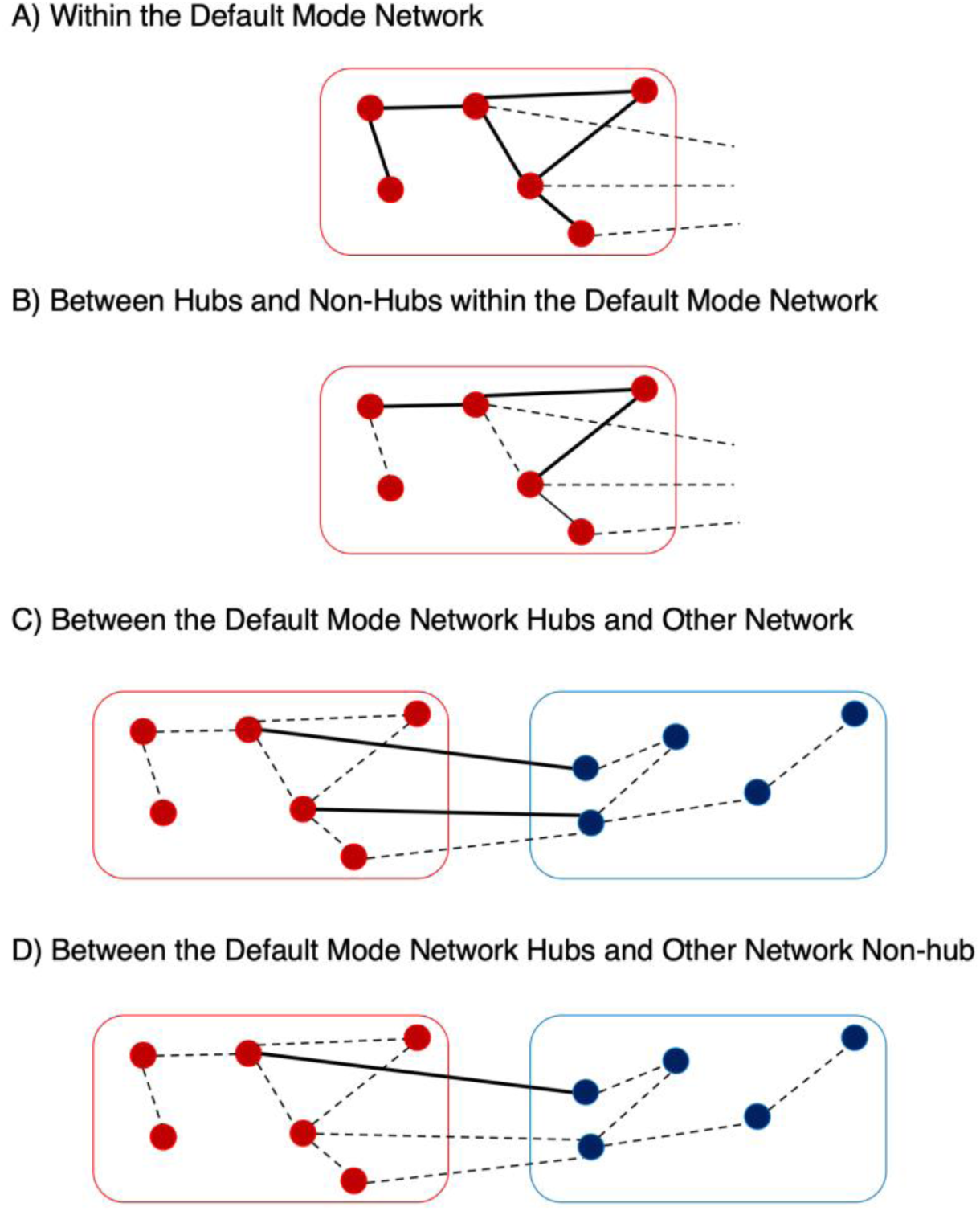
Schematic of Within and Between Default Mode Network Analyses. Nodes are characterized as circles and connections between nodes are characterized by lines. Red nodes represent brain regions within the default mode network (DMN), while blue nodes represent brain regions in a non-DMN network. Hubs are represented as nodes with three or more connections, while non-hubs are represented as nodes with less than three connections. Solid lines depict connections that would have been included in the example analyses while connections represented as dotted lines would not have been included.

### Within DMN Functional Connectivity

Next, we compared functional connectivity within the DMN at baseline, post-injury, and recovery timepoints. To produce a null distribution of functional connectivity, we also assessed 10,000 permutations of a randomized network with the same number of nodes (k = 58) as the DMN but excluding DMN nodes. Differences in functional connectivity at each timepoint within the DMN were compared to this null distribution. Given that this null distribution was created by using a random selection of nodes, each single permutation does not necessarily share the same characteristics as the DMN. However, this method was selected as a more conservative approach to control for Type I errors. We also compared functional connectivity within DMN hubs, within DMN non-hubs, and between DMN hubs and non-hubs across timepoints.

Hubs and non-hubs were also categorized in the 10,000 permutations of randomized networks. We then compared the functional connectivity within hubs, within non-hubs, and between hubs and non-hubs to obtain a null distribution of hub and non-hub connectivity. Differences in functional connectivity at each timepoint within DMN hubs, within DMN non-hubs and between DMN hubs and non-hubs were compared to the null distribution of randomized networks within randomized hubs, within randomized non-hubs, and between randomized hubs and non-hubs. Significant findings were then related to symptom reporting and cognitive performance to examine the relationship between functional connectivity disruptions within the DMN and clinical presentation.

### Between DMN Functional Connectivity

We assessed functional connectivity between DMN hubs and nodes in the remaining 12 networks across the three timepoints: baseline, post-injury, and recovery. These changes were compared to the connections between the permuted networks and the same nodes in the 12 non-DMN networks. For example, when assessing hubs in the DMN to the visual network, functional connectivity for this analysis was compared to permuted networks that excluded nodes in the DMN and visual network. Significant findings between DMN hubs and networks were then related to symptom reporting and cognitive performance to examine the relationship between functional connectivity disruptions and clinical presentation.

### Statistical Analysis

Statistical analyses of functional connectivity and clinical data were conducted in Matlab (v.R2020) using the *anova* and *fitlme* commands. Analysis of variance for linear mixed-effects models was used to account for the missing clinical data and due to their robustness to violations of distributional assumptions^47^. Models included timepoint (i.e., baseline, post-injury, or recovery) as a fixed effect and the random effect of participant ID. Post-hoc tests across timepoints were analyzed using the *ttest* command in Matlab (v.R2020). To control for multiple comparisons, a false discovery rate (FDR) correction was used. Significant results are reported with adjusted p-values for the within DMN analysis (*p_adj_ < 0.002*), between DMN and other networks (*p_adj_ < 0.013*), symptom clusters (*p_adj_ < 0.004*), and cognitive performance (*p_adj_ < 0.001*).

The relationship between functional connectivity and clinical data, such as symptom reporting and cognitive performance, was evaluated in Matlab (v.R2020) using a linear mixed effect model with the *fitlme* command to understand the interaction between timepoint (i.e., baseline, post-injury, or recovery) and functional connectivity to predict either symptom reporting or cognitive performance. Post-injury clinical and MRI data was compared to baseline and recovery data together. Given the exploratory nature of these comparisons, we did not implement FDR-correction. Post-hoc analyses then compared baseline and recovery separately to understand the relationship to post-injury. In the model, we included the random effect of participant ID as well as time between baseline, post-injury, and recovery data collection to control for within-person effects and the variability in time between baseline assessments and when the concussion occurred.

## Results

### Symptom Reports & Cognitive Assessment

The collegiate student-athletes who participated in this study returned-to-play across an expected recovery trajectory after sustaining their concussion. Student-athletes recovered an average of 11 days (median = 8 days). In addition, 91% of these student athletes recovered within 14 days and all but one athlete (98%) recovered within 28 days, consistent with expected recovery^48^.

Table 2 reports findings for symptom factor and cognitive performance changes across time. Participants’ self-reported total symptoms significantly differed across timepoints (*F*(2, 112) = 22.233, *p* < 0.0001), with more symptoms reported at post-injury than baseline (*t* = −4.499, *p* < 0.0001) and more symptoms reported at post-injury than recovery (*t* = 5.355, *p* < 0.0001). We observed differences across timepoints in affective symptoms (*F*(2, 112) = 5.712, *p* = 0.004), cognitive symptoms (*F*(2, 112) = 17.219, *p* < 0.0001), sleep symptoms (*F*(2, 112) = 14.015, *p* < 0.0001), and somatic symptoms (*F*(2, 112) = 19.047, *p* < 0.0001; Figure 3A).

**Figure 3:**
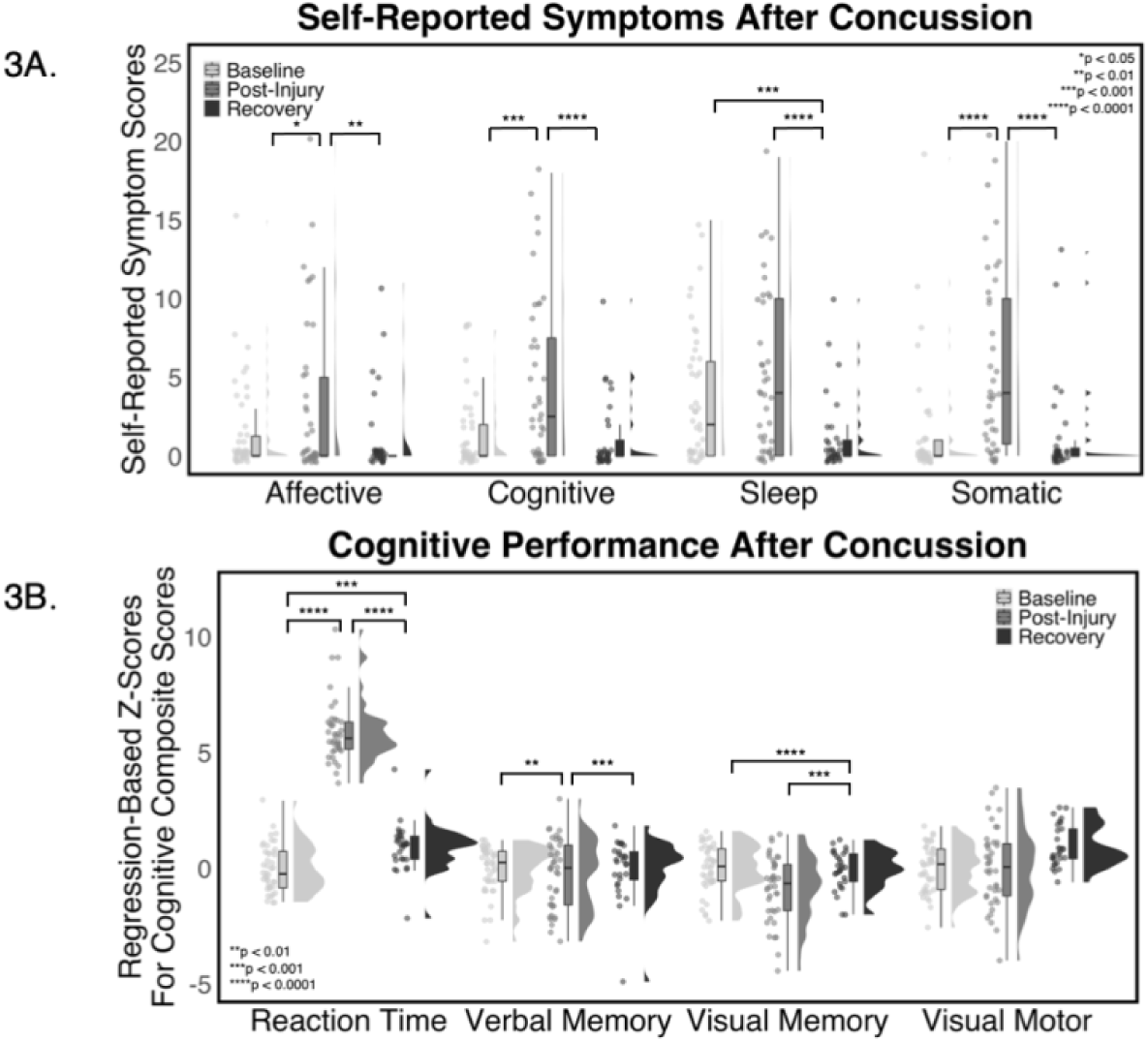
Self-Reported Symptom and Cognitive Performance Changes Change Post-Concussion.

**(A) Changes in the four symptom factors over time.** Affective, cognitive, and somatic symptoms factors increased from baseline to post-injury and then decreased from post-injury to recovery. Less sleep-related symptoms were reported at recovery compared to baseline and post-injury. **(B) Cognitive performance assessed via the ImPACT compositive scores after conversion to regression-based z-scores.** Reaction time slowed from baseline to post-injury and improved from post-injury to recovery, yet performance did not return to baseline levels. Visual memory performance was worse from baseline to post-injury and improved from post-injury to recovery. Visual motor performance was not significantly different between baseline and post-injury but did improve from baseline to recovery. Verbal memory performance did not significantly change across baseline, post-injury, and recovery.

**Table 2.**
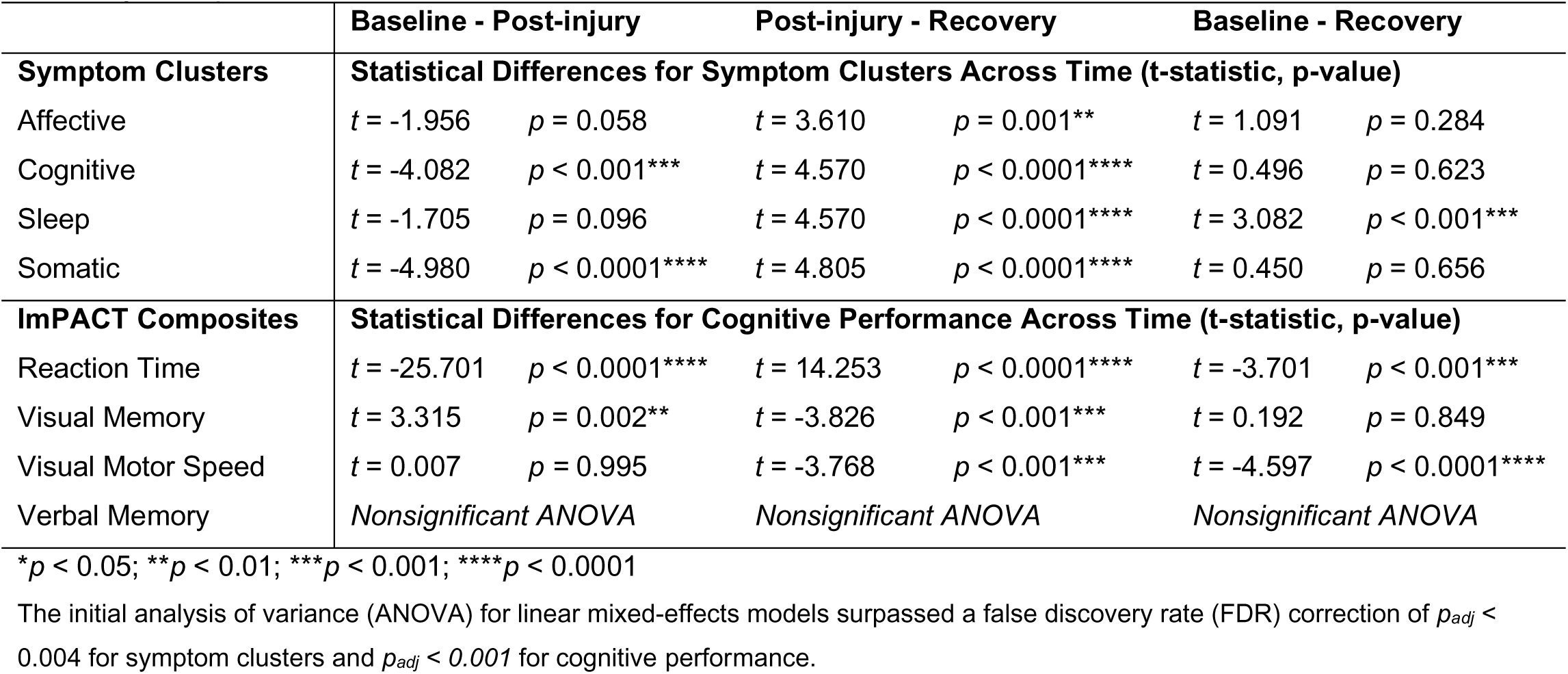
Statical differences for symptom clusters and cognitive performance across baseline, post-injury, and recovery time points.

Student-athlete performance on ImPACT composite scores differed across timepoints for tasks assessing reaction time (*F*(2, 110) = 343.370, *p* < 0.0001), visual memory (*F*(2, 110) = 9.193, *p* < 0.001) and visual motor speed (*F*(2, 110) = 7.254, *p* = 0.001) performance, while verbal memory performance did not significantly change across timepoints (*F*(2, 110) = 0.555, *p* = 0.576; Figure 3B).

### Within DMN Functional Connectivity

Functional connectivity within the DMN did not significantly differ by timepoint (*F*(2, 129) = 8.1, *p* = 0.070), nor were there any significant differences across time for connectivity between DMN hubs (*F*(2, 129) = 9.115, *p* = 0.079) or between DMN non-hubs (*F*(2, 129) = 4.336, *p* = 0.738). However, connectivity between DMN hubs and DMN non-hubs did significantly differ by timepoint (*F*(2, 129) = 13.656, *p* = 0.003; Figure 4), such that connectivity increased from baseline to post-injury (*t* = −4.456, *p* < 0.0001) and decreased from post-injury to recovery (*t* = 3.129, *p* = 0.003), but did not completely return to baseline at recovery (i.e., connectivity at recovery was greater than baseline; *t* = −2.752, *p* = 0.009). A schematic of these analyses is depicted in Figure 2A and Figure 2B. Significant post-hoc analyses are reported in Table 3.

**Figure 4:**
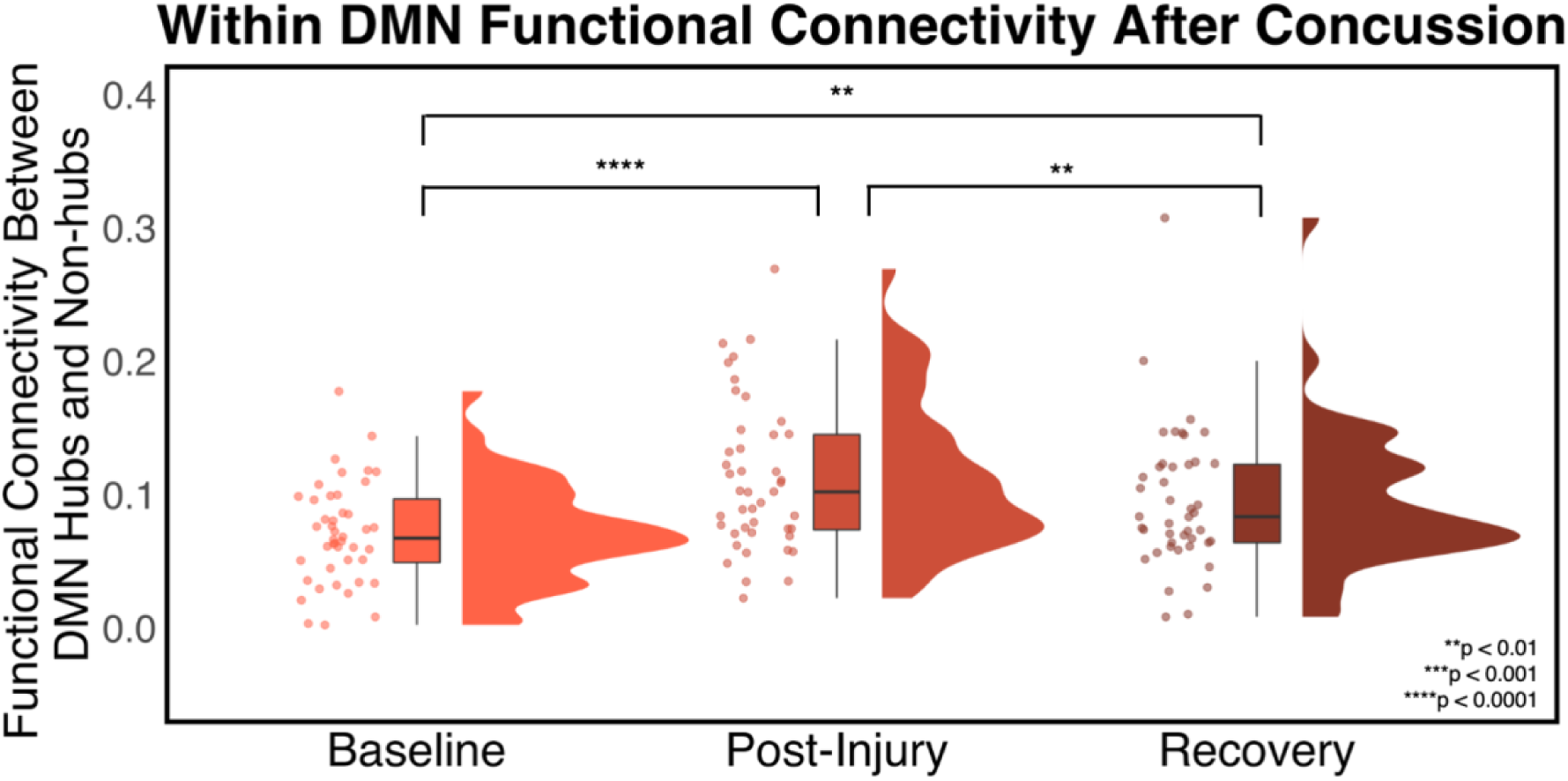
Functional Connectivity Changes Between DMN Hubs and Non-Hubs. Functional connectivity within the DMN between regions identified as hubs and regions identified as non-hubs significantly increased from baseline to post-injury and significantly decreased from post-injury to recovery but did not return to baseline connectivity levels.

**Table 3.**
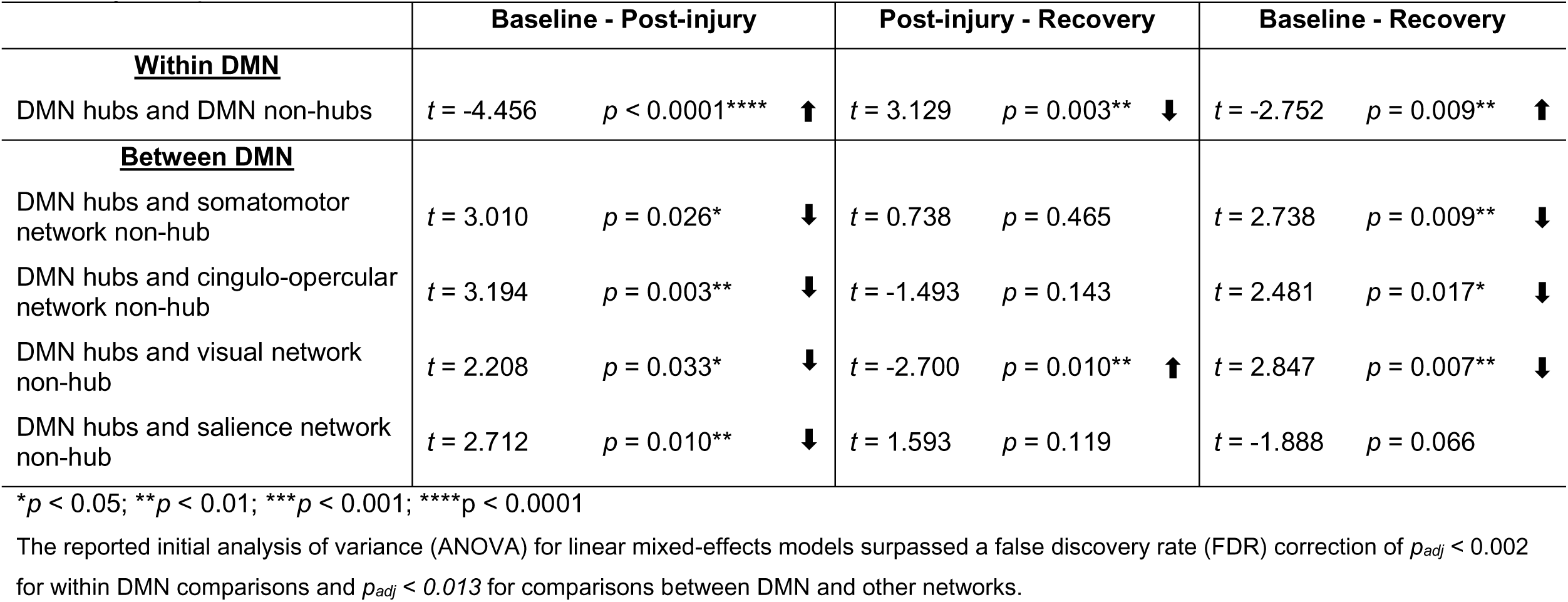
Statistical differences for within and between DMN functional connectivity across baseline, post-injury, and recovery time points.

|  | Baseline - Post-injury |  |  | Post-injury - Recovery |  |  | Baseline - Recovery |  |  |
| --- | --- | --- | --- | --- | --- | --- | --- | --- | --- |
| **<u>Within DMN</u>** |  |  |  |  |  |  |  |  |  |
| DMN hubs and DMN non-hubs | $t = -4.456$ | $p < 0.0001^{****}$ | ↑ | $t = 3.129$ | $p = 0.003^{**}$ | ↓ | $t = -2.752$ | $p = 0.009^{**}$ | ↑ |
| **<u>Between DMN</u>** |  |  |  |  |  |  |  |  |  |
| DMN hubs and somatomotor network non-hub | $t = 3.010$ | $p = 0.026^*$ | ↓ | $t = 0.738$ | $p = 0.465$ | | $t = 2.738$ | $p = 0.009^{**}$ | ↓ |
| DMN hubs and cingulo-opercular network non-hub | $t = 3.194$ | $p = 0.003^{**}$ | ↓ | $t = -1.493$ | $p = 0.143$ | | $t = 2.481$ | $p = 0.017^*$ | ↓ |
| DMN hubs and visual network non-hub | $t = 2.208$ | $p = 0.033^*$ | ↓ | $t = -2.700$ | $p = 0.010^{**}$ | ↑ | $t = 2.847$ | $p = 0.007^{**}$ | ↓ |
| DMN hubs and salience network non-hub | $t = 2.712$ | $p = 0.010^{**}$ | ↓ | $t = 1.593$ | $p = 0.119$ | | $t = -1.888$ | $p = 0.066$ | |

\* $p < 0.05$ ; \*\* $p < 0.01$ ; \*\*\* $p < 0.001$ ; \*\*\*\* $p < 0.0001$
The reported initial analysis of variance (ANOVA) for linear mixed-effects models surpassed a false discovery rate (FDR) correction of $p_{adj} < 0.002$ for within DMN comparisons and $p_{adj} < 0.013$ for comparisons between DMN and other networks.

Table 4 summarizes results from the linear mixed effect models linking this pattern of functional connectivity change with clinical outcomes. Specifically, greater functional connectivity post-injury was observed in participants reporting more cognitive symptoms post-injury resulting in a more positive relationship than the corresponding relationship at baseline and recovery (t = 2.539, p = 0.012), while the relationship at baseline and recovery was not significantly different from one another (t = −0.576, p = 0.566; Figure 5A). Similarly, greater functional connectivity post-injury was observed in participants reporting more somatic symptoms post-injury also resulting in a more positive relationship than the corresponding relationship at baseline and recovery (t = 2.338, p = 0.021), while the relationship at baseline and recovery were not significantly different from one another (t = 0.017, p = 0.987; Figure 5B). Greater functional connectivity post-injury was observed in participants with worse post-injury visual memory performance resulting in a more negative relationship than the corresponding relationship at baseline and recovery (t = −2.793, p = 0.006), while the relationship at baseline and recovery were not significantly different from one another (t = −0.446, p = 0.657; Figure 5C).

**Figure 5:**
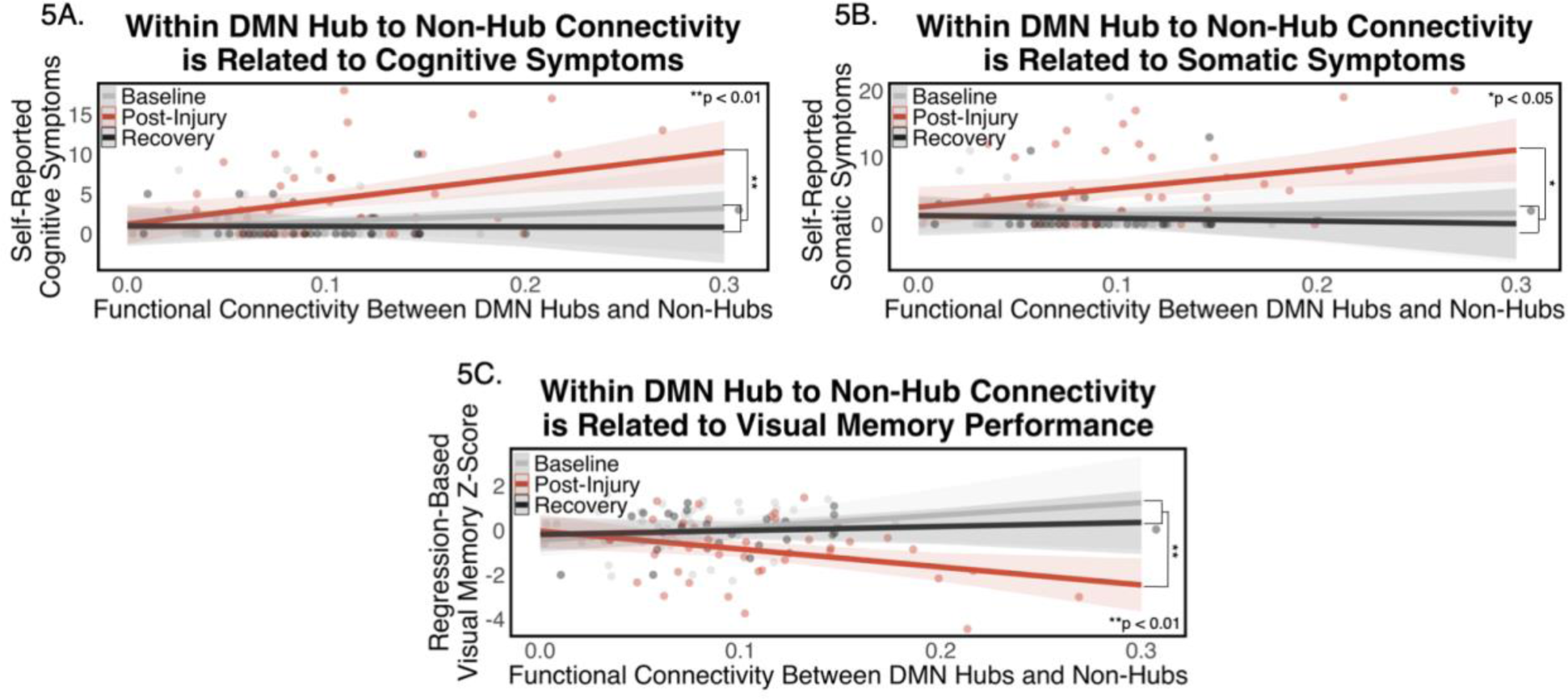
Within DMN Connectivity, Symptom Reporting, and Cognitive Performance. There was a significantly stronger relationship between post-injury functional connectivity between DMN hubs and non-hubs and post-injury **(A)** cognitive symptoms, **(B)** somatic symptoms, and **(C)** visual memory performance than functional connectivity and clinical outcomes at baseline and recovery.

**Table 4.** Significant relationship between post-injury clinical outcomes and functional connectivity between DMN hubs and non-hubs.

| Clinical Outcome | Estimate | Standard Error | T-Statistic | Degree of Freedom | P-value | Confidence Interval | AIC | BIC |
| --- | --- | --- | --- | --- | --- | --- | --- | --- |
| Cognitive Symptoms | 29.213 | 11.505 | 2.539 | 111 | 0.012* | 6.416, 52.011 | 614.1 | 636.1 |
| Somatic Symptoms | 34.285 | 14.663 | 2.338 | 111 | 0.021* | 5.2291, 63.342 | 663.4 | 685.4 |
| Visual Memory | -10.995 | 3.937 | -2.793 | 111 | 0.006** | -18.797, -3.1924 | 348.9 | 370.8 |

\* $p < 0.05$ ; \*\* $p < 0.01$

### Between DMN Functional Connectivity

Functional connectivity from DMN hubs to nodes in the remaining 12 networks were significantly different by timepoint for the cingulo-opercular network (*F*(2, 129) = 8.499, *p* = 0.004), dorsal attention network (*F*(2, 129) = 4.972, *p* = 0.018), salience network (*F*(2, 129) = 6.274, *p* = 0.006), somatomotor network (*F*(2, 129) = 5.677, *p* = 0.019), and visual network (*F*(2, 129) = 13.762, *p* < 0.001). We next focused on connections between hubs in the DMN and non-hubs in the above networks as the connections between hubs and non-hubs within the DMN were specifically altered following concussion. Changes in these connectivity patterns over time were then related to clinical outcomes. A schematic of these analyses is depicted in Figure 2C and Figure 2D.

Functional connectivity differed across timepoint between DMN hubs and non-hubs in the cingulo-opercular network (*F*(2, 129) = 7.605 *p* = 0.002), dorsal attention network (*F*(2, 129) = 3.981, *p* = 0.019), salience network (*F*(2, 129) = 4.980, *p* = 0.026), somatomotor network (*F*(2, 129) = 5.485, *p* = 0.023), and visual network (*F*(2, 129) = 12.861, *p* < 0.0001; Figure 6). Functional connectivity between DMN hubs and cingulo-opercular network non-hubs decreased from baseline to post-injury (*t* = 3.194, *p* = 0.003) and from baseline to recovery (*t* = 2.481, *p* = 0.017). Although functional connectivity between DMN hubs and dorsal attention network non-hubs differed, the pair-wise comparisons did not. Functional connectivity between DMN hubs and the salience network non-hubs only significantly decreased from baseline to post-injury (*t* = 2.712, *p* = 0.010). Functional connectivity between DMN hubs and somatomotor network non-hubs decreased from baseline to post-injury (*t* = 2.310, *p* = 0.026) and from baseline to recovery (*t* = 2.738, *p* = 0.009). Functional connectivity between DMN hubs and visual network non-hubs decreased from baseline to post-injury (*t* = 2.208, *p* = 0.033), increased from post-injury to recovery (*t* = −2.700, *p* = 0.010), and remained decreased from baseline to recovery (*t* = 2.847, *p* = 0.007). These significant post-hoc analyses are reported in Table 3.

**Figure 6:**
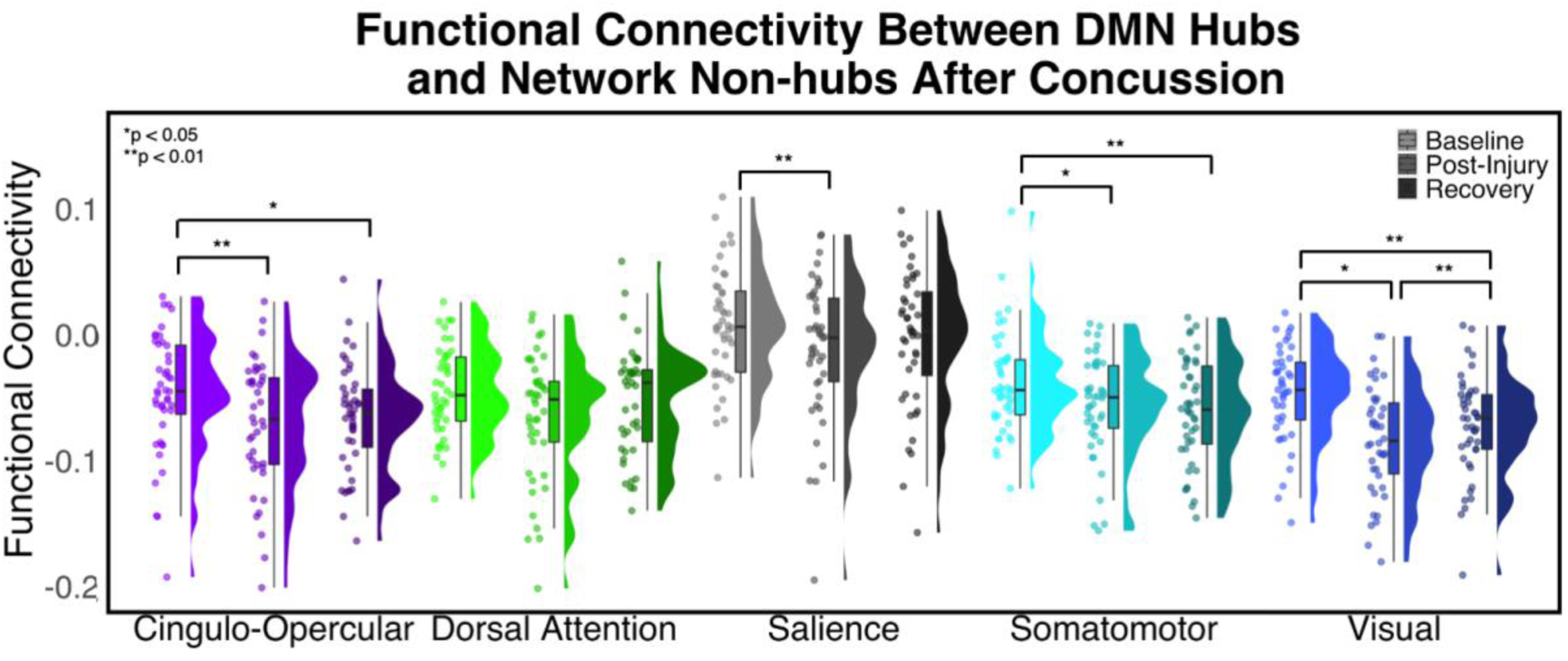
Significant differences in Functional Connectivity Between DMN Hubs and Non-Hubs in Other Networks.

Functional connectivity was significantly different across time between default mode network (DMN) hubs to and non-hubs in five networks. Functional connectivity between DMN hubs and cingulo-opercular network non-hubs decreased from baseline to post-injury and recovery. While functional connectivity between DMN hubs and dorsal attention network non-hubs significantly changed across time, the pair-wise comparisons were not significantly different. Functional connectivity between DMN hubs and salience network non-hubs significantly decreased from baseline to post-injury. Functional connectivity between DMN hubs and somatomotor network non-hubs significantly decreased from baseline to post-injury and recovery. Functional connectivity between DMN hubs and visual network non-hubs significantly decreased from baseline to post-injury, increased from post-injury to recovery, and also decreased from baseline to recovery.

Next, we related this pattern of changes in functional connectivity to symptoms and cognitive performance. The only significant relationship between functional connectivity and symptoms or cognitive performance was with the visual network. Somatic symptoms (Figure 7A) and visual memory performance (Figure 7B) were related to functional connectivity between DMN hubs and visual network non-hubs. Less functional connectivity between DMN hubs and visual network non-hubs at post-injury was observed in participants reporting more post-injury somatic symptoms resulting in a more negative relationship than the corresponding relationship at baseline and recovery (t = −2.466, p = 0.015), while the relationship at baseline and recovery was not significantly different from one another (t = −0.374, p = 0.709). Similarly, for cognitive performance, less functional connectivity at post-injury was observed in participants with worse post-injury visual memory performance resulting in a more positive relationship than the corresponding relationships at baseline and recovery (t = 2.901, p = 0.004), while the relationship at baseline and recovery was not significantly different from one another (t = 0.886, p = 0.378). The significant effects reported here are summarized in Table 5.

**Figure 7:**
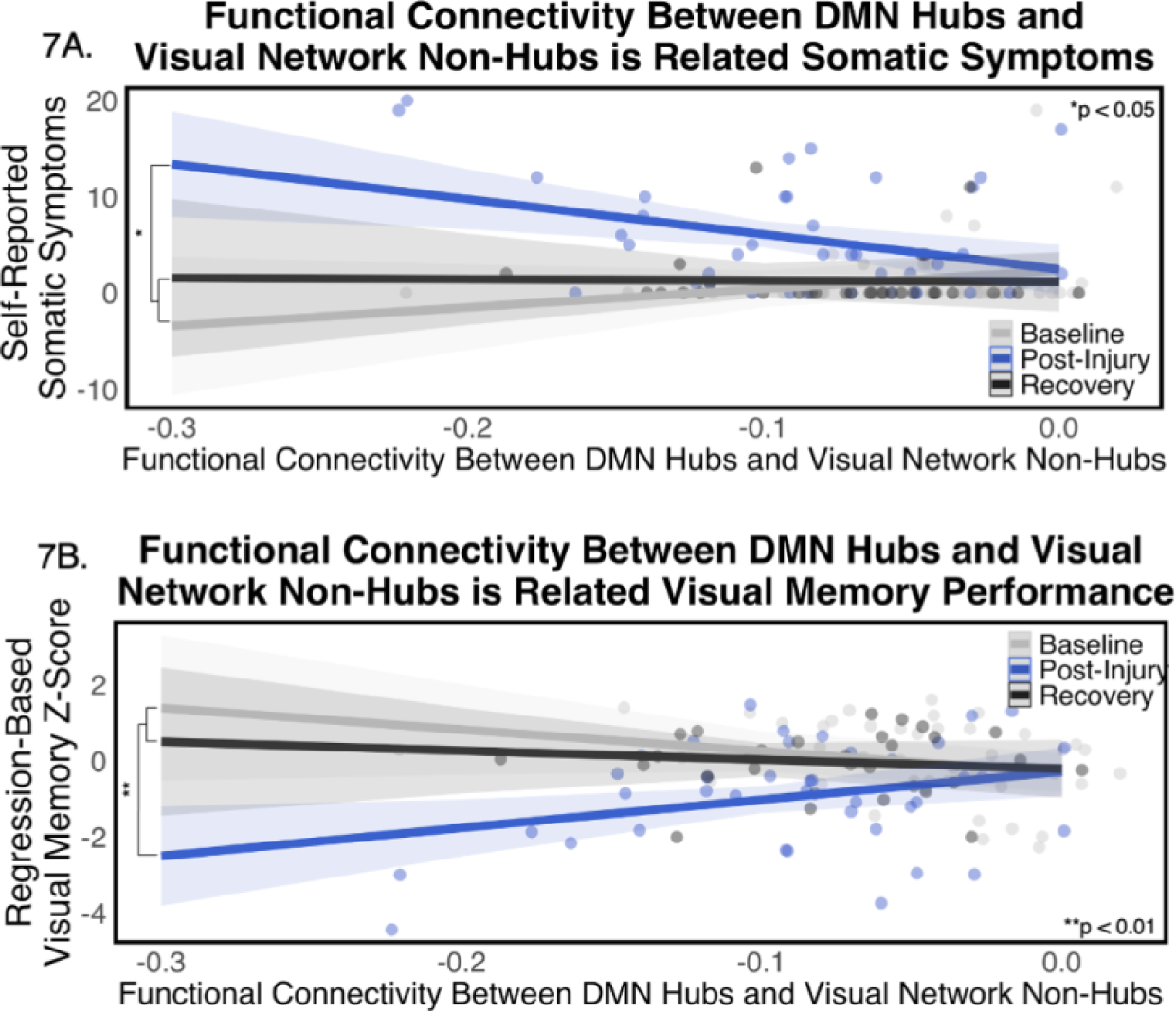
Between DMN and Visual Network Connectivity, Symptom Reporting, and Cognitive Performance (A) Functional connectivity between DMN hubs and visual network non-hubs displayed the only significant between network relationship to clinical outcomes such as symptom reporting and cognitive performance. Specifically, there was a stronger relationship between post-injury functional connectivity and post-injury somatic symptoms than measured at baseline and recovery. **(B)** Similarly, there was a stronger relationship between functional connectivity and visual memory performance at post-injury than at baseline and recovery.

**Table 5.** Significant relationship between post-injury clinical outcomes and functional connectivity between DMN hubs and visual network non-hubs.

| Clinical Outcome | Estimate | Standard Error | T-Statistic | Degree of Freedom | P-value | Confidence Interval | AIC | BIC |
| --- | --- | --- | --- | --- | --- | --- | --- | --- |
| Somatic Symptoms | 29.213 | 11.505 | 2.539 | 111 | 0.012* | -67.176, -7.313 | 663.1 | 685.1 |
| Visual Memory | 11.600 | 3.998 | 2.901 | 111 | 0.004** | 3.6759, 19.524 | 348.5 | 370.4 |

\* $p < 0.05$ ; \*\* $p < 0.01$

## Discussion

Concussion disrupts multiple systems in the body, including functional organization of the brain. The present study examined the effect of concussion in 44 collegiate student-athletes on functional connectivity within and between the default mode network (DMN) and specified the relationship to clinical outcomes after concussion. Uniquely, the present study was able to obtain pre-injury baseline functional connectivity and clinical data in all 44 student-athletes. We found *increased* functional connectivity between hubs and non-hubs within the DMN during the acute period after concussion compared to baseline. At recovery, functional connectivity between DMN hubs and non-hubs *decreased* from post-injury but did not return to baseline levels. In addition, functional connectivity between hubs in the DMN and non-hubs identified in several other brain networks decreased from baseline to post-injury. However, results were variable between networks at recovery with only some networks returning to baseline levels. Clinically, somatic and cognitive symptoms, and visual memory performance were the only measures related to these changes in brain organization post-injury.

### Within DMN Functional Connectivity

We found that post-injury functional connectivity between hubs and non-hubs within the DMN was significantly related to self-reported symptoms and cognitive performance after concussion. Specifically, increased DMN hub and non-hub connectivity was associated with increased cognitive and somatic symptoms after a concussion. While these results are consistent with prior work^23^, we did not find evidence of a relationship with affective symptoms that has been demonstrated (based on depression-related symptoms) in other work^25^. Our findings of functional connectivity between DMN hubs and non-hubs and the relationship to symptomology help expand the understanding of the pathophysiology of clustered symptoms after concussion.

Specifically related to cognition, Li and colleagues (2019) found that hub metrics of frontal regions were associated with a brief measure of cognitive functioning after concussion. Similarly, our findings of DMN functional connectivity, which encompasses frontal regions of the brain, showed a significant relationship with visual memory performance. These results provide additional evidence for the effect of pathophysiological changes in cognitive functioning during the acute stage of concussion recovery. However, it is unclear what role these changes in DMN hubs and non-hubs play specifically in visual memory performance, compared to the other cognitive domains assessed. One explanation relates to the medial prefrontal cortex (mPFC), a central hub of the DMN, and its role in memory^49,50^. Disruption to this hub after concussion may be associated with these acute disruptions to visual memory.

While we identified changes across timepoints between hubs and non-hubs within the DMN, we did not find significant differences by timepoint overall within the DMN. One explanation for the lack of post-concussion functional connectivity changes overall within DMN is because these disruptions may only occur to specialized regions within the network. These more specific changes may primarily impact connectivity to regions that are critical in promoting network efficiency across the brain, such as hubs^51^. Prior research has identified disruption to hubs across the brain in a concussed population^22–25^, but there is lack of consensus in the location of changes. Some studies found that a concussed population had on average a different set of brain regions identified as hubs compared to a non-concussed group and that this set of hubs differed throughout a 6-month recovery period^23,24^. Similarly, we identified that connectivity involving DMN hubs is disrupted after concussion and during recovery. This suggests that concussion results in a reorganization of network efficiency that fluctuates across time; however, it is unclear whether this change represents a positive, negative, or neutral change in brain functioning. Previous studies were unable to compare post-injury changes to pre-injury organization of the brain. A significant contribution of the present study is that we can observe changes in hub connectivity post-injury compared to baseline brain network organization.

### Hyperconnectivity

Considering the severity of brain injury, prior research studying the effects of moderate to severe TBI has identified hyperconnectivity (i.e. increased connectivity) after TBI in both longitudinal studies and in cross-sectional studies compared to healthy participants^12,13,15,52–54^. These studies suggest that after a more severe brain injury, hyperconnectivity may occur as an adaptive response to balance the brain’s network cost and efficiency of brain hubs^12–15^. Specifically related to clinical outcomes, hyperconnectivity of the DMN after moderate to severe TBI has been associated with faster reaction time and better performance on a working memory task^53–55^. A study of mild TBI patients with positive post-injury imaging findings (i.e. evidence of lesions) found that greater DMN connectivity within two weeks of injury was related to better performance on an assessment of attention and processing speed at 6 months post-injury^56^, suggesting a potential early compensatory mechanism for cognitive performance throughout recovery of those with more severe brain injuries such as complicated mild, moderate, or severe TBI. While greater functional connectivity after these more severe injuries has been associated with better cognitive performance, in the present study we found that increased connectivity acutely post-injury, specifically with DMN hubs, was related to *more* cognitive and somatic symptoms, and *worse* visual memory performance. There may still be a potential adaptive mechanism after these less severe injuries, however, the compensation to balance cost and efficiency of brain hubs may occur to a lesser degree in concussion.

Although hyperconnectivity of the DMN has been identified across the TBI severity spectrum, most studies assessing functional connectivity of the DMN after mild TBI have identified variable results of hyperconnectivity and hypoconnectivity post-injury. Many of the studies identifying hypoconnectivity within the DMN compared functional connectivity to healthy control groups and occurred across a range of timeframes post-injury^56–61^. Some of these studies identified a relationship with clinical outcomes including an association between hypoconnectivity, cognitive symptoms^59^ and total symptoms^57,61^, while other studies did not find a significant relationship between DMN connectivity and symptom reporting^58,62^. Additionally, others have found both hypo- and hyperconnectivity between different regions in the DMN compared to healthy controls^62,63^, where hyperconnectivity was related to poorer executive function and hypoconnectivity was associated with increased depression after mTBI^62,63^. Again, the inconsistencies in the literature are likely largely attributed to the lack of pre-injury baseline (i.e., comparing to a control group complicates interpretability given individual differences in cortical organization) and variability in the timeline of assessments.

In other concussion research that explores longitudinal changes in functional connectivity within the DMN, hyperconnectivity was consistently identified during acute recovery^56,60,64^. For example, hyperconnectivity within the DMN was evident within three days of injury but reduced after seven days when neurocognitive assessments had returned to baseline levels^62^. Increased connectivity within the DMN in the earlier stage of recovery was also related to visual and verbal memory performance in athletes^60^. These findings are consistent with the present study and highlight the importance of collecting neuroimaging data in the acute phase of an injury as well as measuring functional connectivity changes over time within participants. These studies, using a longitudinal design to compare functional connectivity changes and clinical outcomes after concussion, clarify previously inconsistent results of hyperconnectivity identified within functional networks. In other words, although hyperconnectivity after moderate to severe TBI is consistently evident when compared to a control group, *mild* TBIs may produce more specific changes in the brain, discussed above, that are more reliant on a within-subjects approach. The understanding of DMN hyperconnectivity and its relationship to symptoms and cognition during recovery described here provides further insight into the brain’s response to pathological changes after injury.

### Between DMN Functional Connectivity

While changes within the DMN are compelling, this study also identified injury-related changes in functional connectivity between DMN hubs and several networks. Connectivity differences between the DMN and visual network were significantly related to somatic symptoms and cognitive performance. Specifically, at post-injury, there was a stronger relationship between functional connectivity between DMN hubs and visual network non-hubs and visual memory performance, as compared to baseline or recovery timepoints. Similar findings have been identified when comparing moderate TBI participants to healthy participants^65^. In addition, another study focused on a hub within the visual network and found functional connectivity was specifically associated with sensitivity to light and noise in a concussed population^23^. Similarly, we also identified a stronger relationship with somatic symptoms, which encompasses light and noise sensitivity, and functional connectivity between DMN and visual network at post-injury. Therefore, future analyses should continue to explore hub and non-hub connectivity of the visual network after concussion to further our understanding of functional connectivity disruptions to this network and their relationship to symptom clusters.

### Changes in Network Segregation Versus Integration

Overall, we observed increased post-injury connectivity within regions of the DMN as well as decreased connectivity between the DMN and multiple networks in the brain. This pattern of results is suggestive of a concussion-related imbalance of within and between network communication. In other words, a healthy brain relies on a balance of processing information globally (e.g., during higher order cognition) through the *integration* of networks as well as relatively specialized (e.g., visual) processing through the *segregation* of networks^19^. This organization allows the brain to efficiently transfer information, when necessary, but also supports functional specialization^20^, thus optimizing the metabolic cost of communicating information throughout the brain^19^. However, when the balance of integration and segregation is disrupted after injury, as evidenced here for concussion, it likely affects the functional output of the brain (as measured by symptom reporting and cognitive performance in this study). Preclinical animal models studying the effects of mild TBI have suggested decreased network integration and increased network segregation for structural connectivity^66^. Similar findings of imbalances of structural connectivity have been identified in children after mild TBI^67^. Further, functional network segregation has also been associated with mild TBI-related fatigue^68^.

Clinically, concussion has been conceptualized as a disorder of multiple network disruptions stemming from changes in processes involving neurometabolic^9^, axonal^69^, cerebrovascular^70^, ocular-motor^71,72^, and vestibular systems^71,72^. The current results provide further evidence of disruption to multiple brain networks during the period following a concussion and lays the foundation to continue relating these network changes to functional outcomes, including specific clustered (e.g., cognitive and somatic) symptoms as well as cognitive difficulties particularly related to the visual system.

### Strengths and Limitations

While these results are informative, this study has several limitations. First, this analysis related functional connectivity findings to clinical outcomes (i.e., self-reported symptoms and cognitive performance) that were collected for clinical purposes. Therefore, we were unable to manage the timing of assessments, resulting in missing data at either post-injury or recovery timepoints for fifteen participants. In addition, baseline clinical data for seventeen participants during our first year of scanning, when the study first began, were collected approximately 1.8 years prior to their MRI scan. However, to reduce the effects of both missing data and variable time between clinical and MRI assessments, we used a linear mixed effects model and included time between data collection as a random effect to help account for the variability in time between baseline assessments and when their concussion occurred. In addition, we did not have a sample of non-concussed student-athletes as a control group. Instead, we used a longitudinal approach that examined brain connectivity in an uninjured state (baseline), and compared it to the acute stage following concussion, and after clearance to return to play. Another strength of the study is that we collected approximately 30 minutes of resting-state fMRI data for each participant at each time point. There is strong evidence that this quantity of data provides reliable estimates of brain network connectivity^73,74^. Lastly, our sample of concussed student-athletes involved 40 men and 4 women. This sample restricted our ability to study gender-specific differences in functional connectivity and recovery, limiting the generalizability of these results to literature discerning differences in recovery for women post-concussion. Future work is necessary to explore these important gender-related differences.

## Conclusion

These results highlight the relationships between acute concussion-related changes to functional connectivity of DMN hubs, visual memory performance, and self-reported cognitive and somatic symptoms, compared to the student-athletes’ own baseline and recovery levels. Specifically, at the time of injury (compared to baseline and recovery), we found hyperconnectivity within regions of the DMN and hypoconnectivity between DMN hubs and distributed networks. These results provide evidence for a disrupted balance of within- and between-network communication, likely impacting network efficiency after concussion. Moreover, these changes in cortical organization were uniquely related to poorer visual memory performance and increased cognitive and somatic symptoms. Notably, some aspects of functional connectivity remained disrupted after clinical recovery, even when symptoms and cognitive performance returned to baseline levels, highlighting the importance of incorporating baseline functional connectivity levels. Understanding pre-injury organization of these athlete’s brain allows us to identify continued change in functional networks which may reflect the brain’s ability to compensate after these changes in network inefficiency.

## Transparency, Rigor, and Reproducibility Statement

This study was not formally registered nor was the analysis plan formally pre-registered. The sample was available from 336 collegiate men’s football and women’s soccer student-athletes who underwent baseline MRI scans starting in June 2018 for football and July 2019 for soccer. Of these participants, 50 experienced a concussion between their baseline scan and October 2021. The same team physician provided a diagnosis for all concussions. Five participants were removed from the analysis for missing either a post-injury or recovery scan and one was removed for excessive motion (> 70% of timepoints were removed due to motion). Thus, 44 participants were included in the analysis. All imaging data were collected using the same 3T Siemens Skyra scanner and preprocessed with the same pipeline. Complete imaging parameters are presented in the methods. Clinical outcomes were collected via ImPACT, a standard tool used for concussion evaluations in the sport setting, and composite scores were converted to regression-based z-scores to account for practice effects and regression to the mean. Fifteen participants were missing clinical outcome data. To account for missing clinical data and to address violations of distributional assumptions, specifically with symptom reports, analysis of variance for linear mixed-effects models were used. Further, a permutation method was selected as a more conservative approach to control for Type I errors when assessing changes in neuroimaging analyses. Replication of the current results is ongoing by the study group. De-identified data from this study are not available in a public archive. Analytic code used to conduct the analyses presented in this study are not available in a public repository. De-identified data from this study will be made available (as allowable according to institutional IRB standards) as well as analytic code by emailing the corresponding author. The authors provided the full content of the manuscript as a preprint as of 03/07/2023.

## Acknowledgements

We wish to acknowledge the help provided by Dr. Lonnie Albers, the University of Nebraska-Lincoln Athletic Medicine team physician, for his involvement in the clinical concussion care of the student-athletes included in this study. We also wish to thank the head football athletic trainer, Mark Mayer, and head soccer athletic trainer, Lisa Loewenstein, for coordinating data collection with the student-athletes at the time of this study. In addition, we wish to acknowledge the help provided by Joanne Murray, Garrett Schwindt, Liv O’Clair, Ruby Basyouni, and Elliot Carlson for aiding in collection of the MRI data. This work was completed using the Holland Computing Center of the University of Nebraska, which receives support from the Nebraska Research Initiative.

## Authorship Contribution Statement

Bouchard conducted analyses, conceptualization, visualization of data, and writing. Higgins provided supervision, resources, collection of clinical data, and contributions to reviewing and editing. Amadon visualized and synthesized data and provided contributions to reviewing and editing. Laing preprocessed data and provided contributions to reviewing and editing. Maerlender provided conceptualization, resources, and contributions to reviewing and editing. Al-Momani preprocessed data and provided contributions to reviewing and editing. Neta provided supervision, methodology, funding, and contributions to reviewing and editing. Savage provided supervision, funding, and contributions to reviewing and editing. Schultz provided supervision, conceptualization, methodology, software and analyses support, and contributions to reviewing and editing.

## References

1. Langlois JA, Rutland-Brown W, Wald MM. The epidemiology and impact of traumatic brain injury: a brief overview. J Head Trauma Rehabil 2006;21(5):375–378; doi: 10.1097/00001199-200609000-00001.

2. Coronado VG, Haileyesus T, Cheng TA, et al. Trends in sports- and recreation-related traumatic brain injuries treated in US emergency departments: the national electronic injury surveillance system-all injury program (NEISS-AIP) 2001-2012. J Head Trauma Rehabil 2015;30(3):185–197; doi: 10.1097/HTR.0000000000000156.

3. Clay MB, Glover KL, Lowe DT. Epidemiology of concussion in sport: a literature review. J Chiropr Med 2013;12(4):230–251; doi: 10.1016/j.jcm.2012.11.005.

4. Pierpoint LA, Collins C. Epidemiology of sport-related concussion. Clin Sport Med 2021;40(1):1–18; doi: 10.1016/j.csm.2020.08.013.

5. Yang J, Comstock RD, Yi H, et al. New and recurrent concussions in high-school athletes before and after traumatic brain injury laws, 2005-2016. Am J Public Health 2017;107(12):1916–1922; doi: 10.2105/AJPH.2017.304056.

6. Sharp DJ, Jenkins PO. Concussion is confusing us all. Pract Neurol 2015;15(3):172–186; doi: 10.1136/practneurol-2015-001087.

7. McCrory P, Meeuwisse W, Dvořák J, et al. Consensus statement on concussion in sport-the 5th international conference on concussion in sport held in Berlin, October 2016. Br J Sports Med 2017;51(11):838–847; doi: 10.1136/bjsports-2017-097699.

8. Silverberg ND, Iverson GL, Arciniegas DB, et al. Expert panel survey to update the American Congress of Rehabilitation medicine definition of mild traumatic brain injury. Arch Phys Med and Rehabil 2021;102(1):76–86; doi: 10.1016/j.apmr.2020.08.022.

9. Giza CC, Hovda DA. The new neurometabolic cascade of concussion. Neurosurgery 2014;75(Supplement 4):S24–S33; doi: 10.1227/NEU.0000000000000505.

10. Chong CD, Schwedt TJ. White matter damage and brain network alterations in concussed patients: a review of recent diffusion tensor imaging and resting-state functional connectivity data. Curr Pain Headache Rep 2015;19(5):12; doi: 10.1007/s11916-015-0485-0.

11. Morelli N, Johnson NF, Kaiser K, et al. Resting state functional connectivity responses post-mild traumatic brain injury: a systematic review. Brain Injury 2021;35(11):1326–1337; doi: 10.1080/02699052.2021.1972339.

12. Hillary FG, Rajtmajer SM, Roman CA, et al. The rich get richer: brain injury elicits hyperconnectivity in core subnetworks. PLoS One 2014;9(8):e104021; doi: 10.1371/journal.pone.0104021.

13. Hillary FG, Roman CA, Venkatesan U, et al. Hyperconnectivity is a fundamental response to neurological disruption. Neuropsychology 2015;29(1):59–75; doi: 10.1037/neu0000110.

14. Hillary FG, Grafman JH. Injured brains and adaptive networks: the benefits and costs of hyperconnectivity. Trends Cogn Sci 2017;21(5):385–401; doi: 10.1016/j.tics.2017.03.003.

15. Roy A, Bernier RA, Wang J, et al. The evolution of cost-efficiency in neural networks during recovery from traumatic brain injury. PLoS One 2017;12(4):e0170541; doi: 10.1371/journal.pone.0170541.

16. Margulies DS, Ghosh SS, Goulas A, et al. Situating the default-mode network along a principal gradient of macroscale cortical organization. Proc Natl Acad Sci U S A 2016;113(44):12574–12579; doi: 10.1073/pnas.1608282113.

17. Schultz DH, Ito T, Solomyak LI, et al. Global connectivity of the fronto-parietal cognitive control network is related to depression symptoms in the general population. Netw Neurosci 2019;3(1):107–123; doi: 10.1162/netn_a_00056.

18. Schultz DH, Cole MW. Integrated brain network architecture supports cognitive task performance. Neuron 2016;92(2):278–279; doi: 10.1016/j.neuron.2016.10.004.

19. Bullmore E, Sporns O. The economy of brain network organization. Nat Rev Neurosci 2012;13(5):336–349; doi: 10.1038/nrn3214.

20. Sporns O. Network attributes for segregation and integration in the human brain. Curr Opin Neurobiol 2013;23(2):162–171; doi: 10.1016/j.conb.2012.11.015.

21. Warren DE, Power JD, Bruss J, et al. Network measures predict neuropsychological outcome after brain injury. Proc Natl Acad Sci USA 2014;111(39):14247–14252; doi: 10.1073/pnas.1322173111.

22. Li F, Lu L, Chen H, et al. Disrupted brain functional hub and causal connectivity in acute mild traumatic brain injury. Aging (Albany NY) 2019;11(22):10684–10696; doi: 10.18632/aging.102484.

23. Bai L, Yin B, Lei S, et al. Reorganized hubs of brain functional networks following acute mild traumatic brain injury. J Neurotrauma 2022;neu.2021.0450; doi: 10.1089/neu.2021.0450.

24. Wang S, Gan S, Yang X, et al. Decoupling of structural and functional connectivity in hubs and cognitive impairment after mild traumatic brain injury. Brain Connect 2021;11(9):745– 758; doi: 10.1089/brain.2020.0852.

25. van der Horn HJ, Liemburg EJ, Scheenen ME, et al. Graph analysis of functional brain networks in patients with mild traumatic brain injury. PLoS ONE 2017;12(1):e0171031; doi: 10.1371/journal.pone.0171031.

26. Buckner RL, Andrews-Hanna JR, Schacter DL. The brain’s default network: anatomy, function, and relevance to disease. Ann N Y Acad Sci 2008;1124(1):1–38; doi: 10.1196/annals.1440.011.

27. Dunkley BT, Urban K, Da Costa L, et al. Default mode network oscillatory coupling is increased following concussion. Front Neurol 2018;9:280; doi: 10.3389/fneur.2018.00280.

28. Sun Y, Wang S, Gan S, et al. Serum neuron-specific enolase levels associated with connectivity alterations in anterior default mode network after mild traumatic brain injury. J Neurotrauma 2021;38(11):1495–1505; doi: 10.1089/neu.2020.7372.

29. Raichle ME. The brain’s default mode network. Annu Rev Neurosci 2015;38:433–447; doi: 10.1146/annurev-neuro-071013-014030.

30. Satpute AB, Lindquist KA. The default mode network’s role in discrete emotion. Trends Cogn Sci 2019;23(10):851–864; doi: 10.1016/j.tics.2019.07.003.

31. Smith V, Mitchell DJ, Duncan J. Role of the default mode network in cognitive transitions. Cereb Cortex 2018;28(10):3685–3696; doi: 10.1093/cercor/bhy167.

32. Puig J, Ellis MJ, Kornelsen J, et al. Magnetic resonance imaging biomarkers of brain connectivity in predicting outcome after mild traumatic brain injury: a systematic review. J Neurotrauma 2020;37(16):1761–1776; doi: 10.1089/neu.2019.6623.

33. Kamins J, Bigler E, Covassin T, et al. What is the physiological time to recovery after concussion? A systematic review. Br J Sports Med 2017;51(12):935–940; doi: 10.1136/bjsports-2016-097464.

34. Bretzin AC, Esopenko C, D’Alonzo BA, et al. Clinical recovery timelines after sport-related concussion in men’s and women’s collegiate sports. J Athl Train 2022;57(7):678–687; doi: 10.4085/601-20.

35. Lovell MR, Collins MW, Podell K, et al. ImPACT: Immediate Post-Concussion Assessment and Cognitive esting. 2000.

36. Lovell MR, Iverson GL, Collins MW, et al. Measurement of symptoms following sports-related concussion: reliability and normative data for the post-concussion scale. Appl Neuropsychol 2006;13(3):166–174; doi: 10.1207/s15324826an1303_4.

37. Merritt VC, Meyer JE, Arnett PA. A novel approach to classifying postconcussion symptoms: the application of a new framework to the Post-Concussion Symptom Scale. J Clin Exp Neuropsychol 2015;37(7):764–775; doi: 10.1080/13803395.2015.1060950.

38. Maerlender AC, Masterson CJ, James TD, et al. Test–retest, retest, and retest: growth curve models of repeat testing with Immediate Post-Concussion Assessment and Cognitive Testing (ImPACT). J Clin Exp Neuropsychol 2016;38(8):869–874; doi: 10.1080/13803395.2016.1168781.

39. Dosenbach NUF, Koller JM, Earl EA, et al. Real-time motion analytics during brain MRI improve data quality and reduce costs. Neuroimage 2017;161:80–93; doi: 10.1016/j.neuroimage.2017.08.025.

40. Gordon EM, Laumann TO, Gilmore AW, et al. Precision functional mapping of individual human brains. Neuron 2017;95(4):791–807.e7; doi: 10.1016/j.neuron.2017.07.011.

41. Talairach J, Tournoux P. Co-Planar Stereotaxic Atlas of the Human Brain: 3-Dimensional Proportional System: An Approach to Cerebral Imaging. Georg Thieme: Stuttgart; New York; 1988.

42. Fischl B. FreeSurfer. Neuroimage 2012;62(2):774–781; doi: 10.1016/j.neuroimage.2012.01.021.

43. Friston KJ, Williams S, Howard R, et al. Movement-related effects in fMRI time-series. Magn Reson Med 1996;35(3):346–355; doi: 10.1002/mrm.1910350312.

44. Power JD, Mitra A, Laumann TO, et al. Methods to detect, characterize, and remove motion artifact in resting state fMRI. Neuroimage 2014;84:320–341; doi: 10.1016/j.neuroimage.2013.08.048.

45. Power JD, Cohen AL, Nelson SM, et al. Functional network organization of the human brain. Neuron 2011;72(4):665–678; doi: 10.1016/j.neuron.2011.09.006.

46. Pedersen M, Omidvarnia A, Shine JM, et al. Reducing the influence of intramodular connectivity in participation coefficient. Netw Neurosci 2020;4(2):416–431; doi: 10.1162/netn_a_00127.

45. Schielzeth H, Dingemanse NJ, Nakagawa S, et al. Robustness of linear mixed-effects models to violations of distributional assumptions. Methods Ecol Evol 2020;11(9):1141– 1152; doi: 10.1111/2041-210X.13434.

48. Broglio SP, McAllister T, Katz BP, et al. The Natural History of Sport-Related Concussion in Collegiate Athletes: Findings from the NCAA-DoD CARE Consortium. Sports Med 2022;52(2):403–415; doi: 10.1007/s40279-021-01541-7.

46. van Kesteren MTR, Ruiter DJ, Fernández G, et al. How schema and novelty augment memory formation. Trends Neurosci 2012;35(4):211–219; doi: 10.1016/j.tins.2012.02.001.

50. Müller NCJ, Dresler M, Janzen G, et al. Medial prefrontal decoupling from the default mode network benefits memory. NeuroImage 2020;210:116543; doi: 10.1016/j.neuroimage.2020.116543.

51. Sharp DJ, Scott G, Leech R. Network dysfunction after traumatic brain injury. Nat Rev Neurol 2014;10(3):156–166; doi: 10.1038/nrneurol.2014.15.

52. Hillary FG, Slocomb J, Hills EC, et al. Changes in resting connectivity during recovery from severe traumatic brain injury. Int J Psychophysiol 2011;82(1):115–123; doi: 10.1016/j.ijpsycho.2011.03.011.

53. Sharp DJ, Beckmann CF, Greenwood R, et al. Default mode network functional and structural connectivity after traumatic brain injury. Brain 2011;134(Pt 8):2233–2247; doi: 10.1093/brain/awr175.

54. Shumskaya E, van Gerven MAJ, Norris DG, et al. Abnormal connectivity in the sensorimotor network predicts attention deficits in traumatic brain injury. Exp Brain Res 2017;235(3):799– 807; doi: 10.1007/s00221-016-4841-z.

52. Bernier RA, Roy A, Venkatesan UM, et al. Dedifferentiation does not account for hyperconnectivity after traumatic brain injury. Front Neurol 2017;8:297; doi: 10.3389/fneur.2017.00297.

53. Palacios EM, Yuh EL, Chang Y-S, et al. Resting-state functional connectivity alterations associated with six-month outcomes in mild traumatic brain injury. J Neurotrauma 2017;34(8):1546–1557; doi: 10.1089/neu.2016.4752.

54. D’Souza MM, Kumar M, Choudhary A, et al. Alterations of connectivity patterns in functional brain networks in patients with mild traumatic brain injury: a longitudinal resting-state functional magnetic resonance imaging study. Neuroradiol J 2020;33(2):186–197; doi: 10.1177/1971400920901706.

55. Iraji A, Benson RR, Welch RD, et al. Resting state functional connectivity in mild traumatic brain injury at the acute stage: independent component and seed-based analyses. J Neurotrauma 2015;32(14):1031–1045; doi: 10.1089/neu.2014.3610.

59. Mayer AR, Mannell MV, Ling J, et al. Functional connectivity in mild traumatic brain injury. Hum Brain Mapp 2011;32(11):1825–1835; doi: 10.1002/hbm.21151.

57. Murdaugh DL, King TZ, Sun B, et al. Longitudinal changes in resting state connectivity and white matter integrity in adolescents with sports-related concussion. J Int Neuropsychol Soc 2018;24(8):781–792; doi: 10.1017/S1355617718000413.

58. Stevens MC, Lovejoy D, Kim J, et al. Multiple resting state network functional connectivity abnormalities in mild traumatic brain injury. Brain Imaging Behav 2012;6(2):293–318; doi: 10.1007/s11682-012-9157-4.

62. Borich M, Babul A-N, Yuan PH, et al. Alterations in resting-state brain networks in concussed adolescent athletes. J Neurotrauma 2015;32(4):265–271; doi: 10.1089/neu.2013.3269.

60. Zhou Y, Milham MP, Lui YW, et al. Default-mode network disruption in mild traumatic brain injury. Radiology 2012;265(3):882–892; doi: 10.1148/radiol.12120748.

64. Zhu DC, Covassin T, Nogle S, et al. A potential biomarker in sports-related concussion: brain functional connectivity alteration of the default-mode network measured with longitudinal resting-state fMRI over thirty days. J Neurotrauma 2015;32(5):327–341; doi: 10.1089/neu.2014.3413.

62. Rahman MRA, Abd Hamid AI, Noh NA, et al. Alteration in the functional organization of the default mode network following closed non-severe traumatic brain injury. Front Neurosci 2022;16:833320; doi: 10.3389/fnins.2022.833320.

63. Meningher I, Bernstein-Eliav M, Rubovitch V, et al. Alterations in network connectivity after traumatic brain injury in mice. J Neurotrauma 2020;37(20):2169–2179; doi: 10.1089/neu.2020.7063.

64. Yuan W, Wade SL, Babcock L. Structural connectivity abnormality in children with acute mild traumatic brain injury using graph theoretical analysis: abnormal structural connectivity in children with acute mTBI. Hum Brain Mapp 2015;36(2):779–792; doi: 10.1002/hbm.22664.

68. Skau S, Johansson B, Kuhn H-G, et al. Segregation over time in functional networks in prefrontal cortex for individuals suffering from pathological fatigue after traumatic brain injury. Front Neurosci 2022;16:972720; doi: 10.3389/fnins.2022.972720.

66. Grossner EC, Mayer AR, Hillary FG. Neuroimaging and sports-related concussion. In: Neuropsychology of sports-related concussion. (Arnett PA. ed) American Psychological Association: Washington; 2019; pp. 119–150; doi: 10.1037/0000114-006.

70. Conder RL, Conder AA. Heart rate variability interventions for concussion and rehabilitation. Front Psychol 2014;5; doi: 10.3389/fpsyg.2014.00890.

68. Kontos AP, Deitrick JM, Collins MW, et al. Review of vestibular and oculomotor screening and concussion rehabilitation. J Athl Train 2017;52(3):256–261; doi: 10.4085/1062-6050-51.11.05.

69. Mucha A, Collins MW, Elbin RJ, et al. A brief Vestibular/Ocular Motor Screening (VOMS) assessment to evaluate concussions: preliminary findings. Am J Sports Med 2014;42(10):2479–2486; doi: 10.1177/0363546514543775.

73. Laumann TO, Gordon EM, Adeyemo B, et al. Functional System and Areal Organization of a Highly Sampled Individual Human Brain. Neuron 2015;87(3):657–670; doi: 10.1016/j.neuron.2015.06.037.

74. Birn RM, Molloy EK, Patriat R, et al. The effect of scan length on the reliability of resting-state fMRI connectivity estimates. Neuroimage 2013;83:550–558; doi: 10.1016/j.neuroimage.2013.05.099.

