## Supplemental Material 1 for "Concussion-related disruptions to hub connectivity in the default mode network are related to symptoms and cognition"

*Within DMN Functional Connectivity for 15% and 35% hub thresholds*

When the threshold of identifying hubs was increased to 35% instead of 25%, functional connectivity between hubs and non-hubs within the DMN remained significant across time ($F(2, 129) = 12.2$, $p = 0.035$) compared to the null distribution of 10,000 permutations. Functional connectivity between DMN hubs and hubs was also significantly different across time ($F(2, 129) = 10.7$, $p = 0.016$), which was not the case when the threshold was set to 25%. However, connectivity between non-hubs and non-hubs within the DMN remained insignificant across time ($F(2, 129) = 4.1$, $p = 0.571$). When the threshold for identifying hubs was decreased to 15% instead of 25%, functional connectivity between hubs and non-hubs within the DMN remained significant across time ($F(2, 129) = 13.4$, $p < 0.0001$). Functional connectivity remained insignificant between hubs and hubs ($F(2, 129) = 3.9$, $p = 0.644$) and between non-hubs and non-hubs within the DMN ($F(2, 129) = 5.4$, $p = 0.606$), which is consistent with results identified with a 25% threshold for hubs. Therefore, when the threshold was either increased or decreased 10%, from the initial chosen 25% threshold, the only significant change identified was functional connectivity between hubs and hubs within the DMN. This was significant when using a 35% threshold and was not when using a 25% threshold.

*Between DMN Functional Connectivity for 15% and 35% hub thresholds*

When the threshold for identifying hubs in the DMN was increased to 35%, functional connectivity from DMN hubs to nodes in the remaining 12 networks were significantly different across time with the auditory network ($F(2, 129) = 5.1$, $p = 0.030$), cingulo-opercular network ($F(2, 129) = 8.2$, $p = 0.002$), dorsal attention network ($F(2, 129) = 6.2$, $p = 0.007$), salience network ($F(2, 129) = 5.5$, $p = 0.009$), somatomotor network ($F(2, 129) = 4.8$, $p = 0.047$), and visual network (F(2, 129) = 11.0, p < 0.0001). These results are consistent with using a 25% threshold. When the threshold for identifying hubs in the remaining 12 networks was increased to 35%, functional connectivity still significantly differed across time between DMN hubs and regions identified as non-hubs in cingulo-opercular network ($F(2, 129) = 8.2$, $p = 0.001$), dorsal attention network ($F(2, 129) = 5.4$, $p = 0.002$), salience network ($F(2, 129) = 5.0$, $p = 0.016$), somatomotor network ($F(2, 129) = 5.1$, $p = 0.033$), and visual network ($F(2, 129) = 10.7$, $p < 0.0001$). Again, these results are consistent with using a 25% threshold. Uniquely, functional connectivity between DMN hubs and auditory non-hubs was significant ($F(2, 129) = 4.1$, $p = 0.036$) with this greater threshold whereas this functional connectivity did not significantly differ when the hub threshold was at 25%.

When the threshold for identifying hubs in the DMN was decreased to 15%, functional connectivity from DMN hubs to nodes in the remaining 12 networks were significantly different across time between DMN hubs and auditory network ($F(2, 129) = 6.0$, $p = 0.018$), cingulo-opercular network ($F(2, 129) = 7.4$, $p = 0.009$), dorsal attention network ($F(2, 129) = 4.8$, $p = 0.024$), salience network ($F(2, 129) = 5.5$, $p = 0.031$), somatomotor network ($F(2, 129) = 7.4$, $p = 0.007$), and visual network (F(2, 129) = 14.3, p < 0.0001). These results are consistent with using a 25% threshold. When the threshold for identifying hubs in the remaining 12 networks was decreased to 15%, functional connectivity still significantly differed across time between DMN hubs and regions identified as non-hubs in cingulo-opercular network ($F(2, 129) = 6.5$, $p = 0.022$), somatomotor network ($F(2, 129) = 6.6$, $p = 0.024$), and visual network ($F(2, 129) = 13.1$, $p < 0.0001$). Uniquely, there was not a significant change across time with this lower threshold between DMN hubs and non-hubs in the dorsal attention network ($F(2, 129) = 3.7$, $p = 0.096$) or salience network ($F(2, 129) = 4.0$, $p = 0.165$).
